# Statistical Investigations of Protein Residue Direct Couplings

**DOI:** 10.1101/334540

**Authors:** Andrew F. Neuwald, Stephen F. Altschul

**Affiliations:** Institute for Genome Sciences and Department of Biochemistry & Molecular Biology, University of Maryland School of Medicine, 670 W. Baltimore Street, HSF III, Room 3053, Baltimore, MD 21201; National Center for Biotechnology Information, National Library of Medicine, National Institutes of Health, Building 38A, 8600 Rockville Pike, Bethesda, MD 20894, USA

**Keywords:** Initial Cluster Analysis, Direct Coupling Analysis, Statistical Significance, Protein sequence-structural analysis

## Abstract

Protein Direct Coupling Analysis (DCA), which predicts residue-residue contacts based on covarying positions within a multiple sequence alignment, has been remarkably effective. This suggests that there is more to learn from sequence correlations than is generally assumed, and calls for deeper investigations into DCA and perhaps into other types of correlations. Here we describe an approach that enables such investigations by measuring, as an estimated p-value, the statistical significance of the association between residue-residue covariance and structural interactions, either internal or homodimeric. Its application to thirty protein superfamilies confirms that DCA scores correlate with 3D pairwise contacts with very high significance. This method also permits quantitative assessment of the relative performance of alternative DCA methods, and of the degree to which they detect direct versus indirect couplings. We illustrate its use to assess, for a given protein, the biological relevance of alternative conformational states, to investigate the possible mechanistic implications of differences between these states, and to characterize subtle aspects of direct couplings. Our analysis indicates that direct pairwise correlations may be largely distinct from correlated patterns associated with functional specialization, and that the joint analysis of both types of correlations can yield greater power. Our approach might be applied effectively to assessing multiple alignment quality, eliminating the need for benchmark alignments. Data, programs, and source code are freely available at http://evaldca.igs.umaryland.edu.

**Author Summary:** The success of Direct Coupling Analysis (DCA) for protein structure prediction suggests that multiple sequence alignments implicitly contain more structural information than had previously been realized, and prompts deeper investigations of the sequence correlations uncovered by either DCA or other approaches. To aid such investigations and thereby broaden the utility of and improve DCA, we describe an approach that measures the statistical significance of the association between DCA and either 3D structure or correlated patterns associated with functional specialization. This approach can be used to obtain better input alignments, and to evaluate the relative performance of DCA methods, their ability to distinguish direct from indirect couplings, and the potential biological relevance and mechanistic implications of alternative conformations and homodimeric interactions.

## Introduction

Contacts among residues largely determine a protein’s three-dimensional structure. Among proteins sharing a common structure, such contacts generally produce correlated substitution patterns between residue pairs. Over evolutionary time substitutions at one residue position often result in compensating substitutions at other positions in order to maintain critical interactions. This allows the prediction of protein structural contacts based upon multiple sequence alignment (MSA) covariance analysis. Early approaches were only partially successful, with a major shortcoming the confounding effect of indirect correlations: When residues at positions *i* and *j* correlate, as do those at positions *j* and *k*, then residues at positions *i* and *k* may also correlate even though they fail to interact directly. Direct Coupling Analysis (DCA) and related methods [1-8] have overcome this problem by disentangling direct correlations from indirect coupling effects. As used here, the term DCA refers to all such approaches. DCA constitutes a major breakthrough in protein structure prediction and is currently being applied successfully on a large scale [9].

DCA programs employ a variety of algorithmic strategies, including *sparse inverse covariance estimation* (PSICOV) [4], *pseudo-likelihood maximum entropy optimization* (EVcouplings-PLM) [5, 6] (CCMpred) [10] and *multivariate Gaussian modeling* (GaussDCA) [11]. DCA methods are evaluated by comparing those residue pairs with the highest direct coupling (DC) scores to residue-to-residue contacts within protein structures. Currently this involves using, for example, ROC curves [11], the Matthews correlation coefficient [12], or F^1^ scores. Such measures are applied in the context of supervised machine learning methods where data points are labeled according to a binary classification scheme; for DCA, those residue pairs that are a specified distance apart within a benchmark structure (e.g., ≤ 5 Å) are labeled as positives and other pairs as negatives. However, due to limited structural information, such labels are often inaccurate. Moreover, there are other reasons to criticize such measures in particular circumstances [13]. In the case of DCA, it is not clear how to interpret such measures when comparing different proteins or distinct structures. To standardize such comparisons, it is desirable to obtain a measure of statistical significance, which also provides insight into how surprised we should be with a given result. As illustrated here, one can use such a measure to determine whether it is better to base DC-scores on a MSA of more closely related proteins rather than on an entire superfamily MSA.

Given a set of structures for a protein superfamily, such a significance measure can help identify those of greatest interest: Direct couplings between pairs of residues presumably are due to selective constraints maintaining functionally important structural interactions. Hence, those protein structures that exhibit the most biologically relevant interactions should achieve the highest level of significance. One could therefore use such a significance measure to select among alternative structural models generated by homology or ab initio structure prediction methods. One may also adapt such a measure to evaluate the degree to which high DC scores are associated with properties other than 3D structural contacts. As illustrated here, for example, one may determine whether those residues most distinctive of a particular protein family are overrepresented among the highest DC-scoring residue pairs.

Here we describe an unsupervised method to estimate, in various contexts, the statistical significance of the correspondence between DCA scores and either protein structural contacts or other protein properties—thereby avoiding the need to label data points. Unlike the current practice of selecting, for analysis, an arbitrary number of the highest scoring pairs (e.g., 0.5 times the query sequence length), our approach determines the optimal number of such pairs automatically based on a statistical criterion, while adjusting automatically for the number of multiple hypotheses tested. Unlike binary classification schemes, our approach takes into account the rank of each residue pair based on both DC scores and 3D distances; hence, it treats the structurally closest residue pairs having high DC scores as of higher biological relevance than such pairs having low scores. By providing a quantitative measure of significance, our approach can detect subtle, yet important features of the data that qualitative measures would fail to distinguish from background noise.

We illustrate this approach by investigating: the relative performance of alternative methods; the biological relevance of alternative structures; subtle structural changes associated with the transition state of Ran GTPase; the contribution of homo-oligomer interfaces to aggregate DC scores; DCA’s dependence on the sequences included in the input MSA; and the correspondence between DCA pairwise correlations and correlated patterns associated with protein functional specialization.

## Statistical Model

Abstractly, we consider being presented with an array of elements ordered by a given primary criterion, and wishing to measure how well it agrees with a secondary criterion that distinguishes and ranks a subset of the elements. More specifically, we seek to identify an optimal initial cluster of elements of the array (defined by a cut), as measured by a relevant p-value. Our approach is based upon Initial Cluster Analysis (ICA) [14], which answers the question: Given a random array of length *L*, containing *D* distinguished elements, represented as ‘1’s, and *L* - *D* ‘0’s, what cut point *X* yields the most surprising initial cluster containing *d* ‘1’s, i.e., those elements up to and including *X*, and what is its probability of occurring? For *L* = 18 and *D* = 7, for example, one such array is “101101<u>1</u>00000010001”, with optimal cut point *X* = 7 (underlined), yielding *d* = 5. Here we note that, in practice, to distinguish elements within our array, we frequently rank all the elements, and distinguish those with rank ≤ *D*. We then might denote our example array as “407205<u>6</u>00000010003” with digits > 0 denoting the ranks of distinguished elements. ICA ignores these ranks when choosing the optimal *X*, whereas we would prefer the *d* distinguished elements to the left of *X* to have higher ranks (i.e., lower numbers) than those to the right. Therefore, we generalize ICA to exploit ranking information by using a ball-in-urn model to calculate a ranking specific p-value *P*_*b*_. In brief, we consider an urn containing *D* balls, of which *d* are colored red. We draw *d* balls from this urn, and calculate the probability *P*_*b*_ that at least *R* of them are red, using the cumulative hypergeometric distribution:

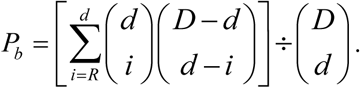

Here, the red balls correspond to those distinguished elements with rank ≤ *d*, and *R* to the number among these elements to the left of the cutoff. (Note that we use *d* both as the number of balls colored red, and as the number of balls drawn from the urn.) A low value of *P*_*b*_ is reported for a cut with a surprising number, among those distinguished elements to its left, having rank ≤ *d*.

Before it corrects for optimizing over all possible cuts, ICA can be understood as calculating a p-value *P*_*a*_ for finding *d* distinguished elements to the left of a cut. Because the calculation of *P*_*a*_ ignores ranking information, it will be independent of *P*_*b*_, and these two p-values may therefore be combined to yield a joint p-value *P*_*J*_ [15-18] using the formula

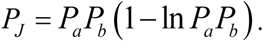

Low values of *P*_*J*_ may arise from low values of *P*_*a*_, or *P*_*b*_, or of both. The p-values *P* we report in this paper correspond to *P*_*J*_, after it has been corrected for optimization over the multiple cut points considered, as described in [14]. One may wish to optimize as well over various values of *D*, but in the current application larger values of *D* are then almost always preferred, due to the indirect couplings considered below. We therefore choose a fixed *D*, based upon a maximum allowed 3D distance within a reference structure.

To apply the theory above to the question of how well DCA methods actually uncover direct contacts within proteins, we proceed as follows. Given an MSA, a method to calculate DCA scores for all column pairs, and a reference structure corresponding to one of the sequences in the MSA, we consider only those pairs of MSA columns separated by ≥ *m* intervening positions within the reference sequence, with *m* = 5 by default. Ordering these column pairs by descending DCA score yields our array of elements, of length *L*. To investigate how well these DCA scores correspond to actual 3D distances, we distinguish those *D* elements whose 3D distance, per the reference structure, is ≤ *r* Å, with *r* = 5 by default. Among the distinguished elements, a column pair having a smaller reference distance receives a higher rank.

### Implementation and availability

We implemented these algorithms and statistical models in C++ as the STARC (<u>S</u>tatistical <u>T</u>ool for <u>A</u>nalysis of <u>R</u>esidue <u>C</u>ouplings) program, which, along with the source code, is freely available at http://evaldca.igs.umaryland.edu.

## Simulations

Here and below, for a calculated theoretical p-value *P* we define a corresponding score as *S* = −log_10_ *P*. Our theory should yield accurate p-values and scores for randomly generated, or shuffled arrays. However, in the present application many column pairs within an MSA are interrelated (e.g., {*i*,*j*}, {*j*,*k*} and {*i*,*k*}), possibly affecting their DCA scores as well as the corresponding distances derived from a structure. To test whether computed p-values remain valid given such interrelationships, we generated a set of 100,000 random p-values as follows: (1) Randomly permute the residues within each aligned column of a glycerol-3-phosphate acyltransferase MSA (cited in Table 1 below) to eliminate residue correlations among columns. (2) Using CCMpred [10], create an ordered array of DCA scores. (3) Compute *P* using the structure 1k30A and a distance cutoff of 5 Å. We define *Ŝ*, as a function of *S*, to be −log10 of the proportion of random scores that are observed to be greater than or equal to *S*. If our p-value calculations are accurate, *Ŝ* should equal *S* to within stochastic error. In **Figure 1** we plot, for *S* from 2 to 5, the *Ŝ* obtained from 100,000 random p-values obtained for arrays generated from permuted MSA columns, as described above. For comparison, we plot as well the *Ŝ* obtained from an equivalent number of shuffled arrays. The straight, solid line represents the agreement of *Ŝ* with theory, and dashed curves represent error ranges of two standard deviations. As can be seen, within stochastic error, *Ŝ* agrees with theory for the shuffled arrays. (Because we can generate p-values rapidly for shuffled arrays, we have confirmed the accuracy of *Ŝ* in this case for *S* ≤ 8; data not shown.) However, for permuted multiple alignment columns, the correlations among array positions yields values of *Ŝ* that are systematically small for *S* > 2.5. Thus, computed values of *S* > 2.5 overestimate significance (i.e. underestimate true p-values) to some degree, but remain valid for comparative analyses, such as those below.

**Table 1.** *S* for thirty superfamilies using residue pairwise 3D distances ≤ 5 Å and a minimum of 5 intervening residues.

| Query | Resolution<br>(Å) | Length | Description | Number of<br>sequences | $S$ for method:<sup>a</sup> | | | |
| --- | --- | --- | --- | --- | --- | --- | --- | --- |
|  |  |  |  |  | CCM<sup>b</sup> | EVC<sup>c</sup> | GSF<sup>d</sup> | PCV<sup>e</sup> |
| 1ayaA | 2.05 | 101 | tyrosine phosphatase SH2 domain | 12,208 | 55 | 54 | **60** | 42 |
| 1b5oA | 2.20 | 382 | aspartate aminotransferase | 105,741 | **892** | 810 | 782 | 609 |
| 1el3A | 1.70 | 315 | aldose reductase | 67,824 | **614** | 588 | 514 | 504 |
| 1k30A | 1.90 | 234 | glycerol-3-phosphate acyltransferase | 12,225 | 111 | **114** | 91 | 78 |
| 1b23P | 2.60 | 184 | elongation factor Tu | 78,839 | **168** | 135 | 111 | 101 |
| 1olzA | 2.0 | 481 | Sema4D | 5,453 | 190 | **245** | 187 | 97 |
| 1wznA | 1.90 | 155 | SAM-dependent methyltransferase | 146,217 | **173** | 156 | 161 | 156 |
| 1z0kC | 1.92 | 164 | Rab4 GTPase | 64,211 | **212** | 181 | 205 | 179 |
| 1zp9A | 2.00 | 258 | Rio1 serine kinase | 24,076 | 105 | **110** | 90 | 86 |
| 2b61A | 1.65 | 357 | homoserine transacetylase | 47,508 | **299** | 290 | 3 | 279 |
| 3ex7H | 2.30 | 241 | DEAD-box ATPase eIF4AIII | 98,478 | **259** | 239 | 175 | 158 |
| 4ag9A | 1.76 | 165 | glucosamine-6-phosphate acetylase | 107,738 | 172 | 174 | **177** | 169 |
| 5dfiA | 1.63 | 318 | apurinic-apyrimidinic endonuclease | 36,297 | 298 | **317** | 245 | 193 |
| 5hf7A | 1.54 | 227 | thymine DNA glycosylase | 7,588 | **128** | 126 | 73 | 67 |
| 5m4pA | 2.30 | 164 | pyruvate dehydrogenase kinase | 1,651 | 33 | 34 | **36** | 23 |
| 1ijxA | 1.90 | 127 | cysteine-rich domain of sfrp-3 | 3,224 | 24 | **25** | 19 | 19 |
| 2nrlA | 0.91 | 147 | myoglobin | 9,514 | **100** | 95 | 70 | 60 |
| 4cmlA | 2.30 | 313 | Inpp5b | 4,724 | 247 | 247 | **270** | 179 |
| 1jw9B | 1.70 | 249 | molybdopterin synthase MoeB | 23,170 | **340** | 318 | 278 | 245 |
| 3fhkF | 2.30 | 147 | disulfide isomerase | 1,042 | 59 | 64 | **65** | 49 |
| 3h7uA | 1.25 | 335 | plant stress-response enzyme Akr4c9 | 67,652 | **598** | 573 | 512 | 490 |
| 1g9rA | 2.00 | 311 | galactosyltransferase LgtC | 10,575 | **283** | 264 | 280 | 211 |
| 4em8A | 1.95 | 148 | ribose 5-phosphate isomerase B | 7,217 | **185** | 181 | 161 | 147 |
| 1i6mA | 1.72 | 328 | tryptophanyl-tRNA synthetase | 20,731 | **327** | 312 | 198 | 168 |
| 3f11A | 0.95 | 252 | oxidoreductase, Ycik | 99,991 | **460** | 448 | 437 | 404 |
| 1jr3A | 2.2 | 373 | bacterial DNA clamp loader $\gamma$ subunit | 24,739 | **386** | 373 | 264 | 248 |
| 1nnlA | 1.53 | 225 | human phosphoserine phosphatase | 130,332 | **138** | 132 | 112 | 92 |
| 1frwA | 1.75 | 194 | E. coli MobA | 79,445 | 192 | 177 | **214** | 170 |
| 1bqbA | 1.72 | 301 | Aureolysin metalloproteinase | 5,289 | **341** | 336 | 253 | 194 |
| 2ovdA | 1.8 | 182 | human complement protein C8γ | 6,874 | 62 | **74** | 51 | 41 |
| average: |  |  |  |  | **248** | 240 | 203 | 182 |

<sup>a</sup>Hydrogen atoms were added using the reduce program [31], except for 3f11A for which hydrogens were already present in the pdb coordinate file. <sup>b</sup>CCMpred. <sup>c</sup>EV coupling pseudo-likelihood maximization. <sup>d</sup>GaussDCA using the Frobenius norm ranking. <sup>e</sup>PSICOV version 2.4 using the recommended $-p$ and $-d$ 0.03 options.

**Figure 1.**
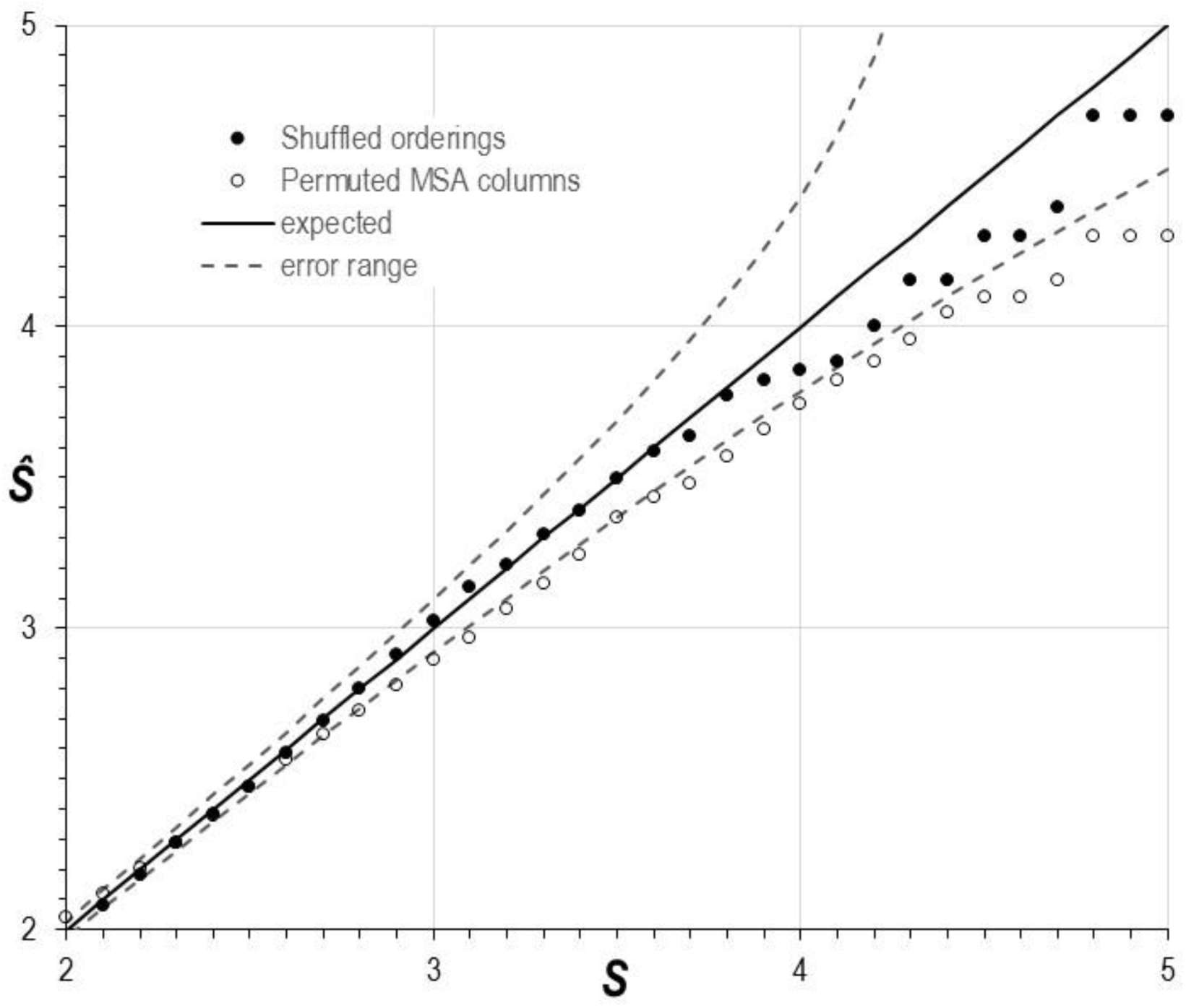
Empirical values of *Ŝ* as a function of *S* yielded by randomly shuffled 100,000 DCA arrays, and by 100,000 DCA arrays derived from MSAs where the residues within each column were randomly permuted. The solid line represents agreement of *Ŝ* with *S*, and the dashed curves represent an error range of two standard deviations.

## Application: Comparisons among DCA methods

We ran the STARC program on the output from four DCA programs, EVcouplings [5, 6], GaussDCA [11], PSICOV [4], and CCMpred [10], each applied to thirty protein domain MSAs with reference 3D contacts ≤ 5 Å (**Table 1**). We scored each run as *S* = −log_10_ (*P*). For a given MSA, better performing programs should generate more significant results, and thus higher scores.

### Comparisons among DCA methods

To evaluate the relative performance of various DCA methods we applied the 2-tailed Wilcoxon signed-rank test [19] to the data in Table 1, after first normalizing each score through division by the total number of residue pairs for its input MSA; the resulting normalized scores approximately follow a Gaussian distribution (see Materials and Methods). Since this test is based on thirty pairs of scores, the sum of the Wilcoxon signed ranks tend to follow a Gaussian distribution. Consequently, we calculated a Z-value for each pair of methods and the corresponding a two-tailed p-value (**Table 2**). This ranked CCMpred as performing marginally better than EVcouplings (*p* = 0.01); EVcouplings better than GaussDCA (*p* = 0.004); and GaussDCA better than PSICOV (*p* = 2×10−6). For individual MSAs, the contribution of *P*_*b*_to *P* varied, for CCMpred, from insignificant to highly significant (e.g., *P*_*b*_ = 6.3×10^−17^ for 3h7uA) with a geometric mean of *P*_*b*_ = 4.6×10^−7^ The superior performance of CCMpred and EV-couplings is not surprising, as both are based on pseudo-likelihood maximization (PLM), which was first introduced as GREMLIN [20] and which was later shown [21, 22] to be more accurate than newer, faster methods such as PSICOV [4].

**Table 2.**
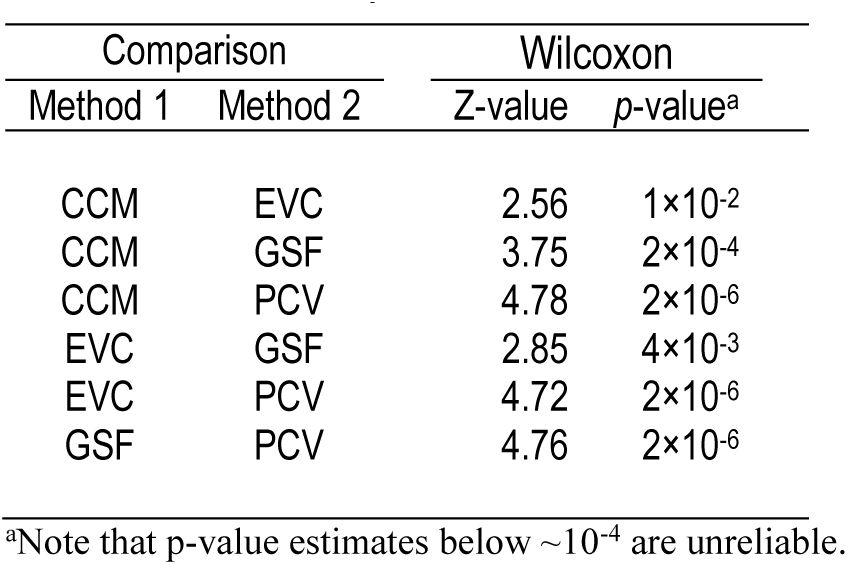
Wilcoxon Signed Rank 2-tailed tests for the 30 analyses in Table 1.

### Indirect couplings

Ideally, as their name indicates, DCA scores should correspond to *direct* correlations between pairs of columns in a MSA. This implies that each DCA pairwise score is statistically independent of every other pairwise score—an assumption upon which our statistical model is based. Consequently, if a DCA method generates output inconsistent with this assumption, by picking up indirect couplings, our approach will identify these as statistically surprising as well. To determine whether this occurs, one can look for significant p-values (i.e., high *S*) arising from pairs of residues distant in the 3D structure. Ideally, in the absence of indirect couplings, *P*s generated by distant pairs alone should not be significant. Note, however, that high *S* for large distances may be due in part to pairs directly coupled in an alternative conformation, or indirectly coupled via functional interactions mediated by other molecules or by a homo-oligomeric interface.

In **Figure 2** we present bar plots for *S* averaged over the thirty superfamilies of Table 1 for residue pairs defined as “discriminating” based exclusively on various distance ranges. (Note that we discarded from the DCA array all pairs corresponding to 3D distances below each specified range.) The high values of *S* we obtained for distant pairs suggests that all four methods are detecting couplings well beyond a residue-to-residue distance of 5 Å—EVcouplings more so than the other methods. For example, in the 2-3 Å range, *S* for CCMpred is significantly higher on average than for EVcouplings (Z-value = 3.57; p = 4×10^−4^), but in the 7-8 Å and 9-10 Å ranges, *S* for EVcouplings is significantly higher (Z=2.89, p = 0.004 and Z=3.24, p = 0.001, respectively).

**Figure 2.**
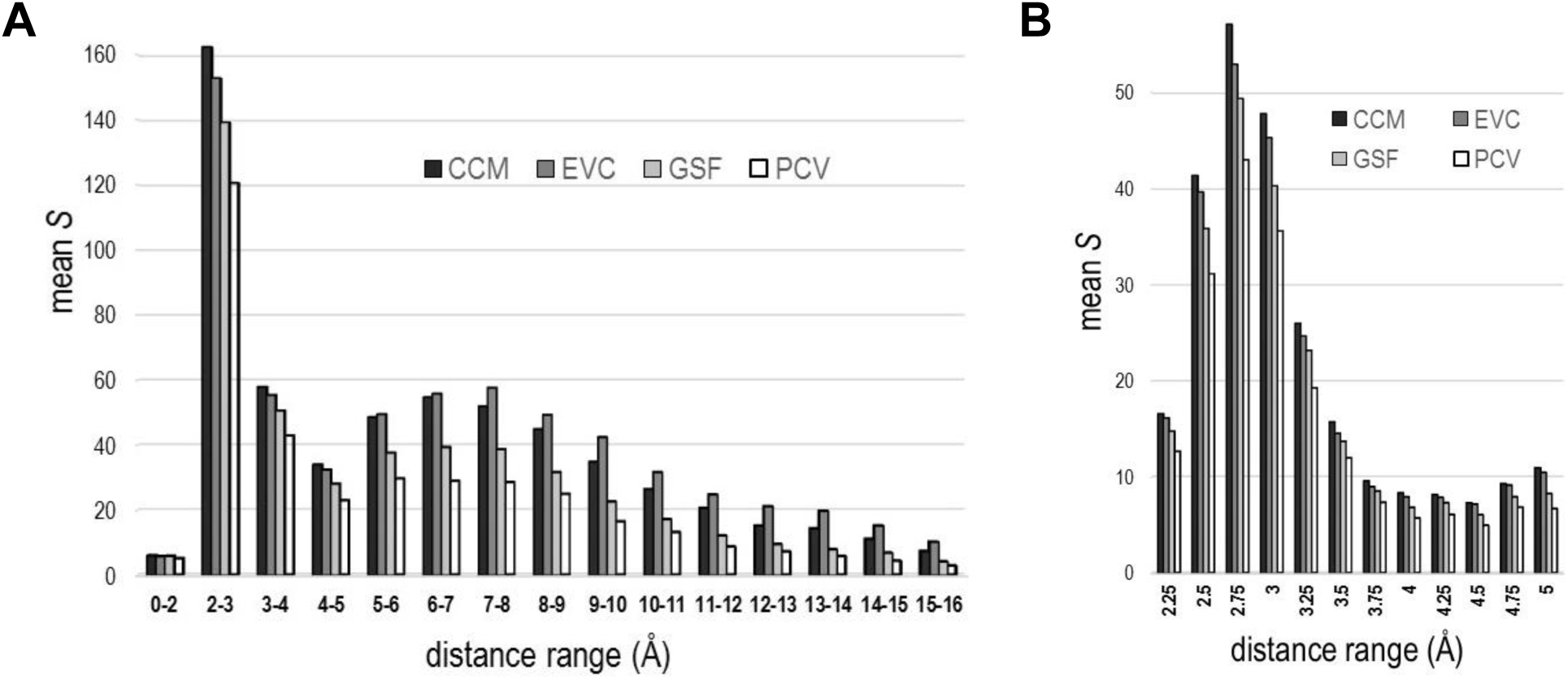
*S* as a function of 3D distance ranges defining distinguished residue pairs. See discussion in text. **A**. Scores obtained for distance ranges spanning zero to 16 Å. Column pairs corresponding to residue-to-residue distances below the indicated range were excluded from the analysis. **B**. Detailed plot of the span 2 to 5 Å. Each distance range covers 0.25 Å and is labeled by its upper limit.

## Application: Quantifying a structure’s biological relevance

We have studied, through the score *S*, the correspondence between a multiple alignment’s DCA scores and the pairwise distances implied by the structure for a particular sequence in the alignment. However, to calculate *S*, there are typically many structures to choose among, and these may differ in important particulars. Recent studies [23-28] have demonstrated that high DC scoring pairs that are distant in certain benchmark 3D structures may come into contact within alternative conformations or across homo-oligomer interfaces, and have thereby provided insight into protein biophysical and dynamic properties. Other studies [29, 30] have combined DCA with correlation analyses involving larger groups of structurally and/or functionally correlated residues, thereby generating further insight. Here we illustrate the application of our method to these sorts of studies.

To the degree to which DCA scores capture the pairwise correlations imposed by the functional requirements common to a protein family, we expect the *S* yielded by a particular structure to reflect the degree to which that structure exhibits critical interactions characteristic of the family. In other words, *S* may measure the degree to which a specific structural conformation is biologically relevant. To investigate this, we consider three cases - human Ran GTPase, Gna1 *N*-acetyltransferase from *C. elegans*, and the bacterial (*E. coli*) clamp loader complex. Using available structures for each of these, we add hydrogen atoms using the Reduce program [31] to better discriminate among residue-to-residue contact distances. A previous DCA analysis [27] found that the heavy atom distance distribution for directly coupled residue pairs exhibited local maxima at 2.8 Å and 3.7 Å, which were interpreted as corresponding to the *donor-acceptor distance* of hydrogen bonds and to hydrophobic interactions, respectively. Here we choose to focus on hydrogen bond interactions. Since our analyses explicitly model hydrogen atoms, we calculate *S* using a maximum structural distance of 2.6 Å, which, based on the sum of the van der Waals radii for hydrogen plus either nitrogen or oxygen [32], corresponds to an upper bound on the *hydrogen-acceptor distance* of hydrogen bonds.

### Ran GTPase

Ran GTPase is required for the translocation of proteins and RNA through the nuclear pore complex. Ran exists in both GDP-and GTP-bound forms. The nucleotide exchange factor RCC1 converts Ran-GDP into Ran-GTP. Ran-mediated hydrolysis of GTP to GDP, which is believed to drive transport of cargo from the nucleus into the cytoplasm, involves the combined action of Ran GTPase activating protein (RanGAP), which activates Ran’s intrinsic GTPase activity, and of the Ran-binding proteins RanBP1 [33].

Two crystal structures of the Ran-RanBP1-RanGAP ternary complex are available [34]: one in the ground state (i.e., bound to a non-hydrolysable GTP analog) and another in a transition-state mimic. For each crystal structure, the unit cell contains four tertiary complexes whose Ran subunits are labeled as chains A, D, J and G. Each chain yields an *S* for each of the two structures, as shown in **Table 3**, and, on average, the *S* for the transition-state exceeds that for the ground state by > 22 based on the R^4^ family MSA described below. (Note that, for Ran, we find no correspondence between *S* and crystal structure resolution, as shown in **Figure 3**.) This average difference in *S*, corresponding to greater than 22 orders of magnitude in *P*, indicates that the transition state forms more functionally relevant interactions than does the ground state. A detailed investigation of the transition state interactions absent from the ground state may provide insight into this key step in Ran-mediated nuclear transport. We investigate this possibility in **Figure 4A** by showing those residues participating in pairs that, for all four Ran subunits within the crystal structure unit cell: (1) are < 2.6 Å apart (and thus before the cutoff *X* in the DCA array) for the transition state, but not for the ground state; and (2) are closer by ≥ ⅓ Å in the transition state than in the ground state. These residues appear to form allosteric pathways between Ran’s active site and its sites of interaction with RanBP1 and with RanGAP. The latter site includes a salt bridge, between Lys130 of Ran and Asp225 of RanGAP, that contributes to the stimulation of GTP hydrolysis by RanGAP [34]. In contrast, residues that participate in pairwise interactions that are relatively stable among diverse conformational forms occur in regions adjacent to these putative pathways (**Figure 4B**). Notably, Phe90, which forms a stabilizing interaction with Gly121 in the guanine binding loop [35], and Val14 are the only residues that (based on our criteria) participate in both transition-state-specific interactions and stable interactions, and therefore may function as pivot points. This analysis illustrates how one may use our approach to investigate structural changes of potential functional relevance.

**Table 3.** *S* for Ran GTPase in the transition state complex^a^ and in the corresponding ground state complex^b^. Scores are based on CCMpred DCA with *L* = 12,090, on a minimum of 5 intervening <u>residues, and on a m</u>aximum distance of 2.6 Å.

| Input MSA: | number of<br>seqs | transition state *S* |  |  |  |  | ground state *S* |  |  |  |  | Δ*S* |
| --- | --- | --- | --- | --- | --- | --- | --- | --- | --- | --- | --- | --- |
|  |  | A<sup>c</sup> | D | J | G | avg | A | D | J | G | avg | avg |
| GTPase superfamily | 274,681 | 55.7 | 55.6 | 55.1 | 55.7 | 55.5 | 44.2 | 42.7 | 48.1 | 43 | 44.5 | 11 |
| R<sup>4</sup> family<sup>d</sup> | 27,571 | 96.9 | 98.7 | 96.8 | 93.7 | 96.5 | 72.7 | 73.9 | 75.6 | 72.3 | 73.6 | 22.9 |
| Ran subfamily | 507 | 5.5 | 5.6 | 5.1 | 5.4 | 5.4 | 6.7 | 6.8 | 6.7 | 6.7 | 6.7 | -1.3 |

<sup>a</sup>Ran-GDP-AIFx-RanBP1-RanGAP; pdb: 1k5g; 3.1 Å.
<sup>b</sup>Ran-GppNHp-RanBP1-RanGAP; pdb: 1k5d; 2.7 Å.
<sup>c</sup>The letters A, D, J and G correspond to the chain designations for each of four Ran subunits within the crystal structure unit cell.
<sup>d</sup>The R<sup>4</sup> family is composed of multiple subfamilies; this includes Rab, Rho, Ras and Ran GTPases.

**Figure 3.**
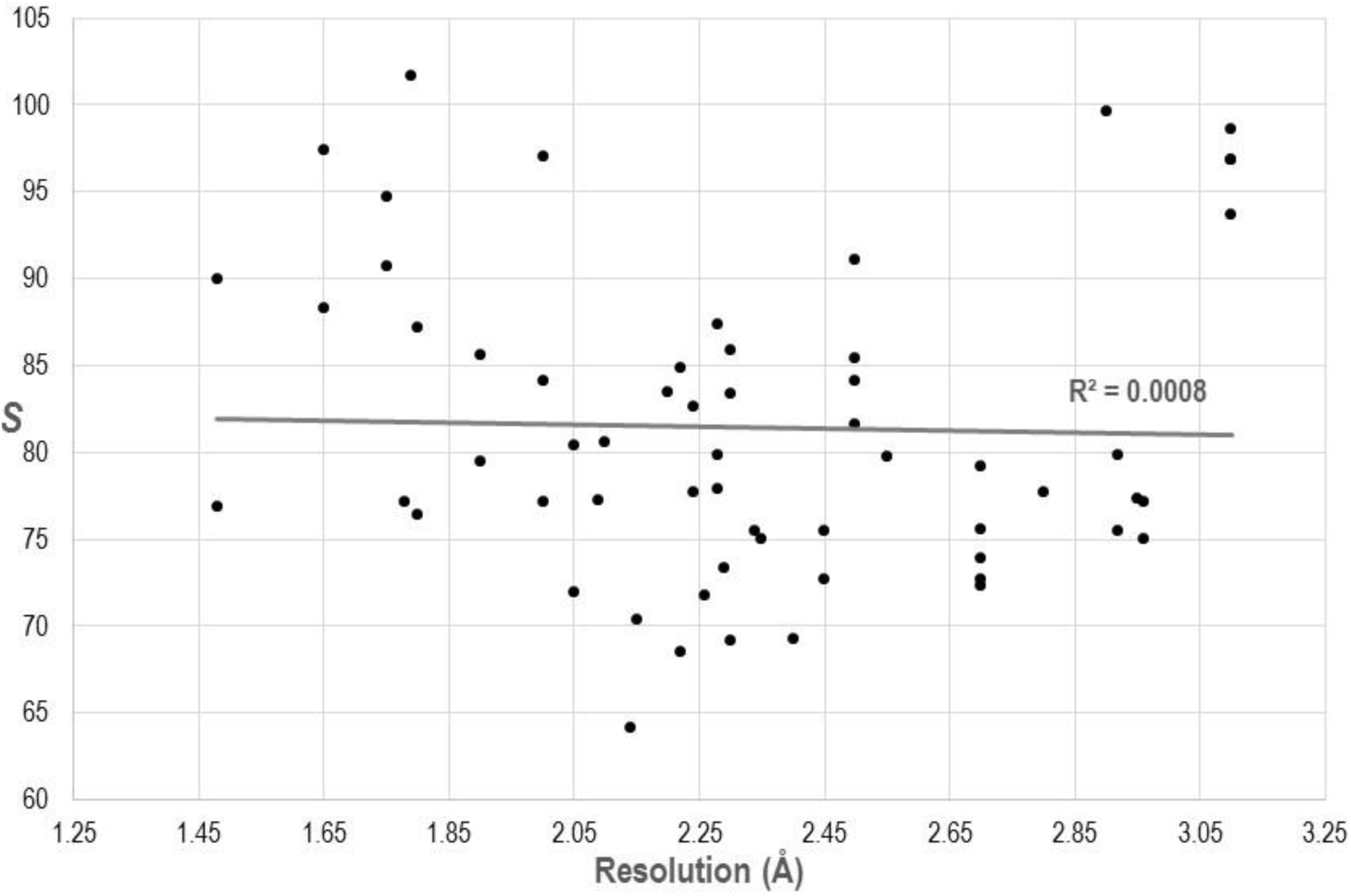
Regression analysis of scores for 60 Ran GTPase structures versus their crystal structure resolutions. The coefficient of determination, *R*^2^, is near zero, indicating that crystal structure resolution fails to explain the *S*’s variability around its mean. The same R^4^ family MSA and parameters were used here as for the analyses in Table 3.

**Figure 4.**
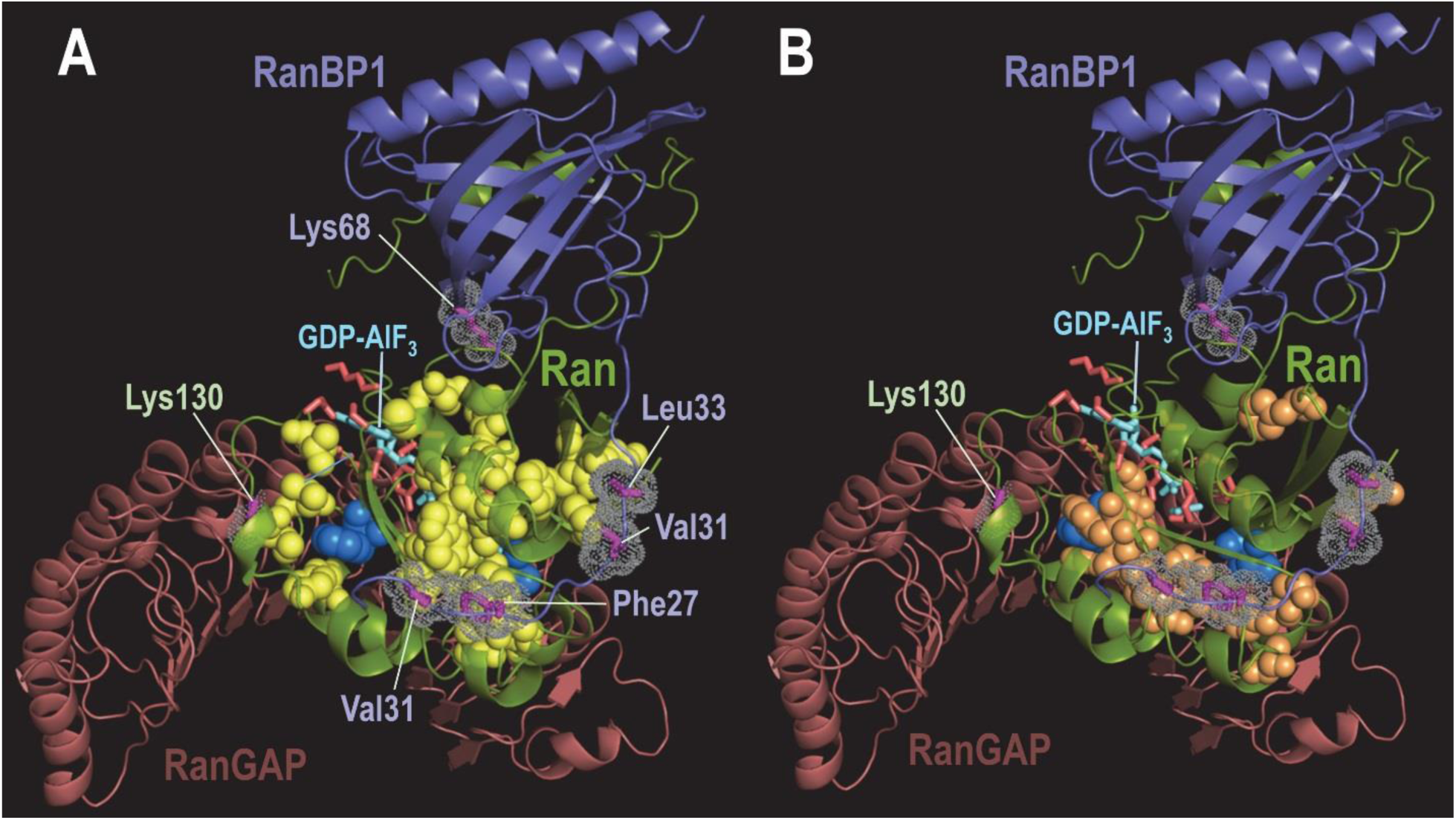
Residues in Ran involved in interacting pairs within the transition state structure (pdb: 1k5g) [34]. Sidechains of residues in RanBP1 contacting Ran are labeled in (A) and shown in magenta with dot clouds. The sidechain of Ran Lys130, which plays a role in the stimulation of GTP hydrolysis by RanGAP [34], is indicated. Sidechains of Ran’s catalytic (active site) residues and the GTP transition state analog are represented as red and cyan sticks, respectively. A PyMOL session file corresponding to this figure is available at our website. **A**. Sidechains of residue pairs contributing to the higher *S* for Ran in the transition state versus the ground state (pdb: 1k5d; structure not shown). These residues are represented as yellow spheres, except for the pivot point residues Phe90 and Val14, which are shown as bright blue spheres, and for two of the catalytic residues (Thr24 and Thr42), which are shown as red sticks. **B**. Ran residues forming pairs whose interactions remain stable over diverse conformational forms are shown as orange and bright blue spheres. The conformational forms include the Ran-RanBP1-RanGAP transition (pdb: 1k5g) and ground (pdb: 1k5d) states; Ran bound to its exchange factor, RCC1 (pdb: 1i2m); Ran bound to GDP (pdb: 3gj0); Ran bound to NtF1 and GDP (pdb: 1a2k); and Ran bound to RanBP1 and CRM1 (pdb: 4hb2).

### Higher S for a Ran subgroup of P-loop GTPases

To examine the dependence of *S* on the sequences included in the input MSA, we used the Bayesian Partitioning with Pattern Selection (BPPS) program [36] to classify the aligned sequences into three nested sets consisting of all P-loop GTPases, of Rab, Rho, Ras and Ran (termed R^4^) GTPases, and of Ran GTPases. We calculated *S* from the DCA scores for each of these MSAs based on the Ran subunit of the Ran-RanBP1-RanGAP transition and ground state structures (**Table 3**). On average, *S* based on the R^4^ family exceeded *S* based on the GTPase superfamily by 41 and 29 for the transition and ground states, respectively. This suggests that proteins within the R^4^ subgroup share pairwise constraints and mechanistic similarities that other P-loop GTPases lack.

### Complementarity of DCA and BPPS analyses of Ran

Like DCA, BPPS identifies correlations among MSA columns, but unlike DCA it focuses on detecting family-specific sequence patterns associated with functional specialization rather than on pairwise correlations. There may be some overlap between the patterns of correlation detected by the two approaches, but this overlaps is often fairly weak. For illustrative purposes, we consider the DCA array used for the analysis of the R^4^ family in Table 3. As shown in **Figure S1A**, when pairs of positions separated by ≤ 5 Å are distinguished, the optimal initial cluster, highlighted in yellow, is highly significant (*S* = 231); 65% of the pairs in this cluster are distinguished, and 53% of all distinguished pairs are in the cluster. These high percentages reflect DCA’s success in detecting directly interacting residues. BPPS defines the R^4^ family by recognizing positions having distinctive residue patterns, and, for comparison to Figure S1A, we distinguish in **Figure S1B** the elements of the DCA array corresponding to pairs of these positions. Again there is a significant (*S* = 6.2) initial cluster, highlighted in yellow. However, only 6% of the pairs in this cluster are distinguished, and only 16% of all the distinguished pairs are in the cluster. Thus, while there is a weak tendency for pairs of positions recognized by BPPS as characterizing the R^4^ family to receive high DCA scores, a sizable majority of these pairs do not. In general, DCA and BPPS often recognize correlations of a complementary character. BPPS, in focusing on positions whose residue patterns are distinctive of a particular family, often recognizes correlations among positions on the protein’s surface or far removed spatially, and whose interaction is not direct but rather linked through common function [36].

### Homodimeric Gna1 N-acetyltransferase

For the preceding analysis, we examined spatial contacts only within single protein subunits, whereas correlated mutations are also associated with contacts at homo-oligomer interfaces. To consider such contacts as well, we applied our approach to the homodimeric structure of glucosamine-6-phosphate N-acetyltransferase (Gna1) [37], a GCN5-like *N*-acetyltransferase (GNAT) [38] that transfers an acetyl group from coenzyme A (CoA) to glucosamine-6-phosphate to produce *N*-acetyl-D-glucosamine-6-phosphate (GlcNAc-6P). [In a previous study [39], we found that the residues most characteristic of the GNAT family to which Gna1 belongs are contributed by both subunits to form the active site at the homodimeric interface. This contrasted with the GNAT superfamily’s most characteristic residues, which are remote from the active site.] To study the influence on *S* of including homodimeric interface contacts, either in Gna1 bound to CoA or in Gna1 bound to both CoA and the reaction product GlcNAc-6P, we computed pairwise distances based either solely on contacts internal to each subunit or on both internal and interface contacts. In the latter case, we used the shorter of the two contact distances to rank each residue pair. Our analysis (**Table 4A**) yielded the following observations: (1) Including trans-homodimer contacts significantly increased *S* for the product-bound complex (Δ*S* = 8.3 and 11.6), but decreased *S* for the unbound complex (Δ*S* = −1.6 and −4.0). (2) When only internal contacts were considered, *S* for the product-bound complex failed to increase significantly relative to *S* for the unbound form (Δ*S* = 0.3 and −2.0). (3) In contracts, when trans-homodimer contacts where considered as well, *S* for the product-bound complex increased significantly relative to *S* for the unbound form (Δ*S* = 10.2 and 13.6). This suggests that binding of the product (and, presumably, the substrate) brings into contact residues across the interface to form the active site.

**Table 4.** *S* as a measure of biological relevance for the *N*-acetyltransferase Gna1 bound to either coenzyme A (CoA) (pdb:4ag7; 1.55 Å) or to both CoA and *N*-acetyl-D-glucosamine-6-phosphate (GlcNAc-6P)(pdb: 4ag9; 1.76 Å) and of the overlap between pairs of BPPS pattern residues and the highest DCA scoring pairs.

| **A. *S* based on residue pairwise distances<sup>a</sup>** |  |  |  |  |  |  |
| --- | --- | --- | --- | --- | --- | --- |
| structural state | distance-based *S* |  |  |  |  |  |
|  | A<sup>b</sup> | A:B<sup>c</sup> | Δ*S* | B<sup>b</sup> | B:A<sup>c</sup> | Δ*S* |
| Gna1 + CoA | 49.1 | 47.5 | -1.6 | 49.7 | 45.7 | -4.0 |
| Gna1 + CoA + GlcNAc-6P | 49.4 | 57.7 | 8.3 | 47.7 | 59.3 | 11.6 |
| Δ*S* | 0.3 | 10.2 |  | -2.0 | 13.6 |  |
| **B. *S* based on pairs of BPPS pattern residues<sup>d</sup>** |  |  |  |  |  |  |
| characteristic pattern | pattern-based *S* |  | pattern residue positions<sup>d</sup> |  |  |  |
| GNAT superfamily | 2.2 |  | 153,116,83,118,154,103,31,121,113,24,106,82,108,117,145,149,137,25,148,73,36,122,79 |  |  |  |
| Gna1 family | 27.9 |  | 135,90,92,104,68,95,93,44,141,136,43,102,105,134,40,58,36,101,98,94,35,116,89,54,61 |  |  |  |
| Δ*S* | 25.7 |  |  |  |  |  |

<sup>a</sup>Scores are based on CCMpred DCA, using the corresponding MSA from Table 1, with a maximum distance of 2.6 Å and a minimum of 5 intervening residues.
<sup>b</sup>The letters A and B correspond to the chain designations for the homodimeric subunits; *S* in these columns is based solely on internal contacts.
<sup>c</sup>*S* in these columns is based on internal and interface contacts; the letter to the right of the colon represents the subunit from which trans-homodimer pairwise distances were obtained.
<sup>d</sup>Pattern residues were determined in [36, 39]; positions are ordered by decreasing BPPS score.

Because the homodimeric interface includes many pattern residues characteristic of the Gna1 family [39], we also considered to what extent the DCA and BPPS analyses are complementary (**Table 4B**). Unlike for Ran GTPase, the highest ranked DCA residue pairs correspond, with high significance (*S* = 27.9), to pairs of the 25 highest BPPS-ranked residues characteristic of the Gna1-family. Thus, the degree of complementarity between DCA and BPPS is protein-specific. Note, however, that the overlap between Gna1-family BPPS pairs and either DCA or 3D contacting pairs is far from optimal (**Figure S2**), suggesting that, in this case as well, pairs of the highest ranked BPPS residues are fairly distinct from residue pairs with the highest DCA ranks or with the shortest 3D distances.

### DNA clamp loader complex

To further explore the possible relationship between a structure’s biological relevance and its score *S*, we examined subunits of the bacterial DNA clamp loader complex. This complex forms a spiral-shaped semicircle of two inactive subunits, δ and δ’, and three γ ATPase subunits arranged in the order: δ-γ-γ-γ-δ’. The last two γs and δ’ each functionally interact with the ATP-binding site of the preceding γ subunit. This complex loads a sliding clamp onto primer template DNA. The Ψ protein binds to the clamp loader, thereby coupling it to single-stranded DNA-binding protein. Upon binding to DNA and ATP, Ψ promotes the clamp-loading activity of the complex by stabilizing it in a spiral-shaped conformation consistent with recognition of both RNA and DNA primers [40].

We analyzed two different clamp loader structures: one of the unbound clamp loader complex (pdb_id: 1jr3) [41] and another of the clamp loader bound to primer template DNA and to the Ψ protein and with an analog of ATP bound to each of the γ subunits (pdb_id: 3gli) [40]. First, using the jackhammer program [42], we created one MSA for each of the subunits: δ, γ and δ’, and used CCMpred to generate an ordered DCA array from each MSA. Second, we calculated values of *S* for each array using corresponding structures for the δ, γ and δ’ subunits (**Table 5A**). Note that in the bound form, there are two clamp loader complexes in the unit cell of the crystal structure, yielding two distinct structures for each of the five subunits. The difference Δ*S* between the scores for the bound and unbound forms, shown in Table 5, range from 14 to 57, all highly significant. This conforms to the expectation that the biologically more relevant bound conformation will yield higher *S* than the unbound form, and further illustrates how *S* can be used to evaluate a structure’s biological relevance. However, unlike for Gna1, the inclusion of contacts between adjacent γ subunits (Table 5A) decreases *S*, suggesting that, in this case, homo-oligomer interactions fail to impose detectable constraints. Finally, we explored for clamp loader subunits the putative contributions to direct couplings of hydrogen bond interactions (pairwise distances ≤ 2.6 Å; Table 5A) versus hydrophobic interactions (pairwise distances ≥ 3 Å and ≤ 5 Å; **Table 5B**). This comparison suggests that the biologically relevant clamp loader state favors presumably more geometrically specific hydrogen bond interactions over presumably less specific hydrophobic interactions.

**Table 5.**
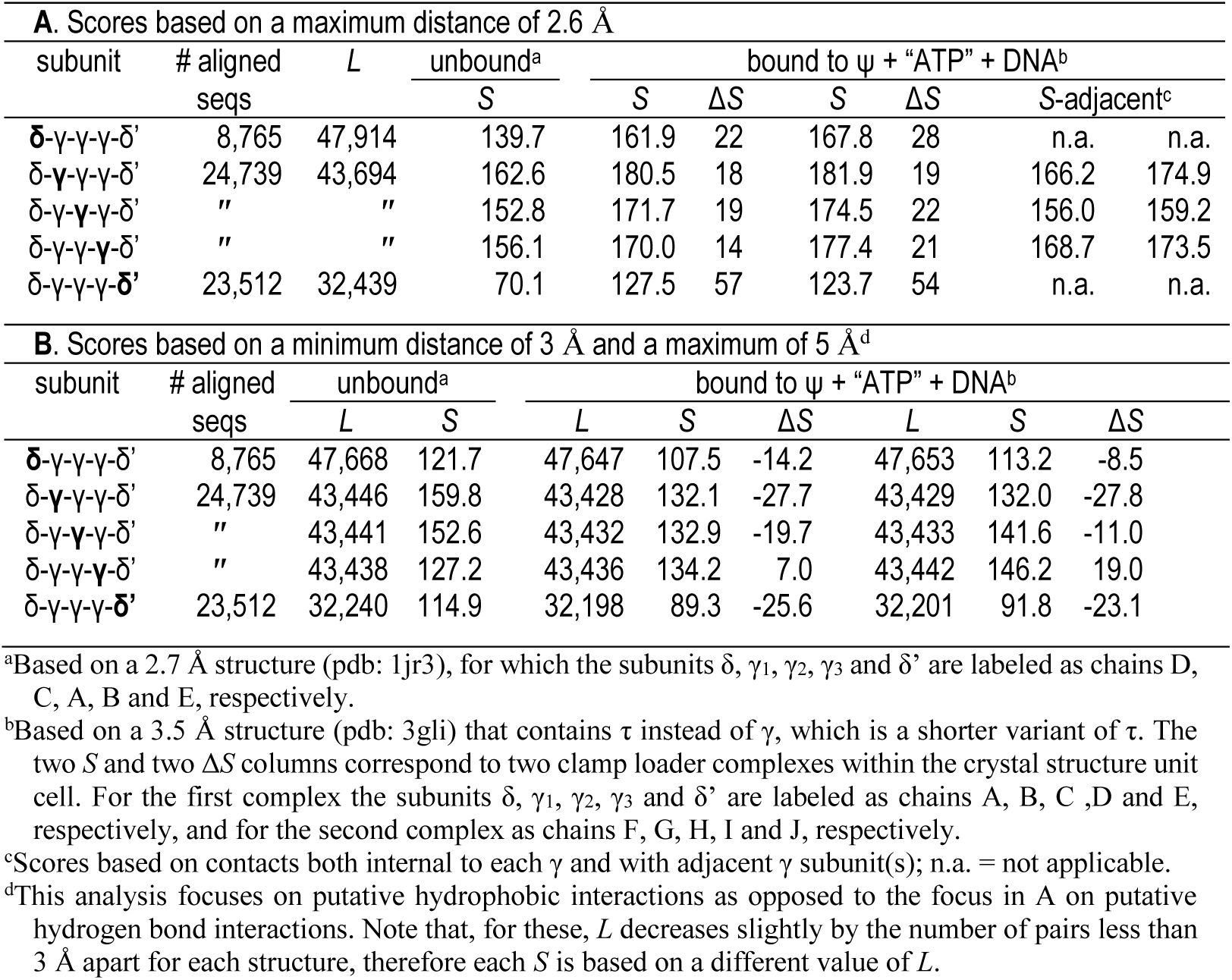
*S* as a measure of structural biological relevance for the bacterial DNA clamp loader complex. Scores are based on CCMpred DCA with a minimum of 5 intervening residues.

## Discussion

*S* scores quantify a DCA method’s ability to detect 3D residue-to-residue contacts. When used in combination with the Wilcoxon signed rank test, they provide a significance measure of the performance of one method versus another and can quantify, as well, a method’s tendency to detect indirect couplings.

We could further develop the STARC statistical model by considering the arrangement of the *d* distinguished pairs before *X*. A pair with a higher DCA score should be more likely than one with a lower score to correspond to a 3D interaction. Ideally, the *d* pairs should thus be arranged in order of decreasing DCA score. To measure how closely a DCA method’s output comes to achieving this configuration, we may proceed as follows: (1) Rank each of the *d* distinguished pairs based on DCA scores, with higher scores receiving lower ranks. (2) Let ∏ be the set of all permutations of the integers [1,d], and for *π* ∈∏ define the score 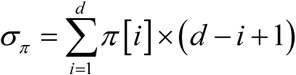. (3) Let ∏_*x*_ be the set of all permutations in ∏ with score *σ*_*π*_ = *x*. (4) Then 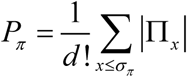 is the probability of obtaining a permutation π with score ≤ *σ* _*π*_. In the absence of an analytic formula, we may compute *P*_*π*_ by exhaustive enumeration, or when this is not tractable,estimate it by Monte Carlo simulation. We may approximate small values of *P*_*π*_ by assuming that it follows a cumulative normal distribution. However, it is unclear whether *P*_*π*_ is independent of *P*_*a*_ and *P*_*b*_, whether the scoring function *σ*_*π*_ is in any sense optimal, and whether there is biological benefit to including this order in our statistical model. We plan to investigate these questions.

An important potential application of our approach, which is beyond the scope of this study, is the evaluation of MSA accuracy without the need for benchmark alignments, which typically contain a relatively small number of sequences and whose accuracy may be uncertain [43]. Our proposed approach would proceed on the assumption that, given available structures, more accurate MSAs will yield higher values of *S*. We are developing this approach, which should benefit from the large amount of sequence data becoming available.

Our analysis of Ran, Gna1 and the DNA clamp loader complex suggests that *S* may be useful for evaluating the biological relevance of alternative structural conformations of the same protein and for characterizing the nature of conformation-specific interactions. Viewing direct couplings as functionally imposed constraints and proteins as molecular machines, *S* may measure the degree to which a particular crystal structure captures a protein in a mechanistically important state. If so, then analyzing in what ways various residue pairs contribution to *S* may provide mechanistic clues. Likewise, comparative analyses among MSAs corresponding to a protein’s subfamily, family and superfamily may provide mechanistic clues regarding functional specialization. Our analysis here also suggests that one may use STARC to search for the most biologically relevant among the very many structures often available for a major protein superfamily.

## Materials and Methods

### Protein structural coordinates

For the thirty STARC analyses in Table 1 we obtained high quality crystal structures from the RCSB Protein Data Bank (PDB) (www.rcsb.org/pdb). The pdb and chain identifiers are given in column 1 of Table 1. Likewise, the coordinates for the Ran, Gna1 and DNA clamp loader analyses were obtained from the PDB; their pdb identifiers are given in Tables 3, 4 and 5, respectively. For all analyses, hydrogen atoms were added using the Reduce program [31], except for the pdb coordinate file for 3F^1^lA in which hydrogens were already present. Hence, residue-to-residue distances are based on any two atoms, including hydrogens, albeit ignoring main chain to main chain interactions. This allows better discrimination among hydrogen bond interactions based on subtle differences in contact distances.

### DCA methods

EVcouplings (EVC) was run over the EVfold website (http://evfold.org) using the pseudo-likelihood maximization (PLM) option with default settings. For each analysis, taking as input the sequence corresponding to the reference structure as the query, EVcouplings uses jackhammer [42] to create a MSA, from which it then computes the direct coupling scores. It returns the jackhmmer alignment in fasta format and the score file. The score file and the corresponding PDB coordinates serve as the input to STARC. We also used the jackhmmer alignment as input to the other programs. The GaussDCA program was run with Frobenius norm ranking (with default parameters); this was done interactively under Julia (www.julialang.org). PSICOV version 2.4 was run using the author recommended –p and –d 0.03 options using as input the jackhmmer alignment after reformatting by the fasta2aln program included with the PSICOV package (http://bioinf.cs.ucl.ac.uk/downloads/PSICOV). CCMpred version 0.3.2 (https://travis-ci.org/soedinglab/CCMpred) was run with default settings using as input the reformatted jackhammer alignment. Note that the output from GaussDCA, CCMpred and PSICOV does not include the query sequence, which, along with the DCA scores, were provided as input to STARC.

### Simulations

For the analysis in Figure 1, we randomly shuffled the DCA arrays using a heapsort routine with randomly generated keys. Likewise, we permuted MSA columns by randomly reordering the residues in each column using heapsort.

### Wilcoxon signed rank test

We evaluated the performance of alternative DCA methods using the Wilcoxon signed-rank test [19], first dividing each *S* by the total number of residue pairs *L*. For CCMpred, EV-couplings and GaussDCA, these normalized scores then approximately follow a Gaussian distribution, as indicated by the Shapiro-Wilk test statistic [44] (p = 0.49, 0.60, and 0.10, respectively). For PSICOV the test score corresponded to p = 0.035, which is slightly below the acceptance threshold of p > 0.05.

### The STARC algorithm

We modified the Initial Cluster Analysis (ICA) algorithm [14] to find the optimal score *S*, as described above. STARC converts PSICOV and GaussDCA formatted DCA score files into EVcouplings format automatically; this requires as input the query sequence in fasta2aln format. We modified the CCMpred source code and recompiled the program to generate PSICOV-formatted output files. Source code for STARC is freely available at: http://evaldca.igs.umaryland.edu/.

## Acknowledgements

We thank L. Aravind and Natarajan Kannan for helpful comments. SFA is supported by the intramural research program of the National Institutes of Health, National Library of Medicine. AFN is supported by the NIH-NIGMS (grant number R01GM125878). The funders had no role in study design, data collection and analysis, decision to publish, or preparation of the manuscript.

---

**Table S1.**
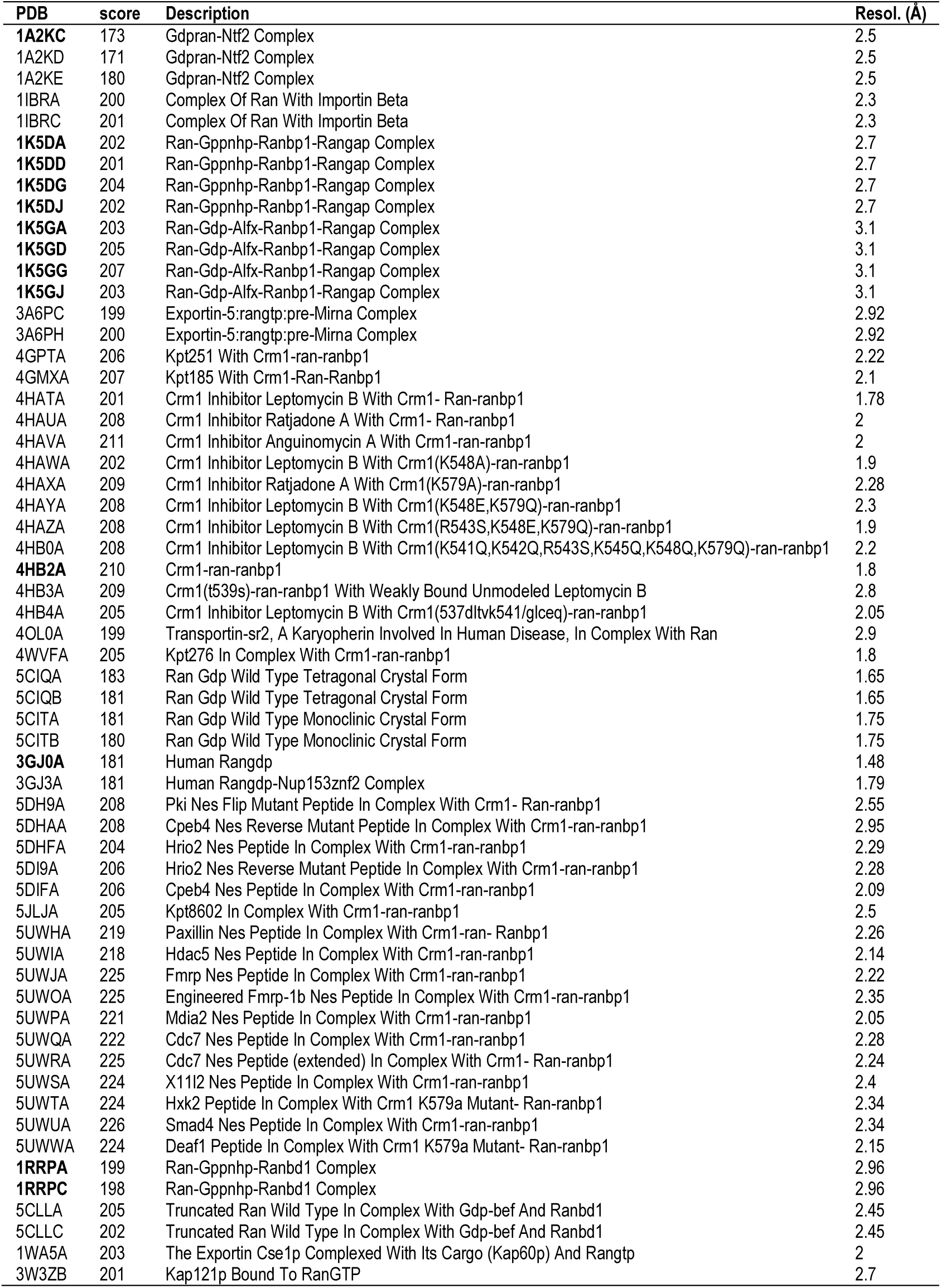
Protein structural coordinates used for the Ran GTPase analysis in Figure 3.

**Figure S1.**
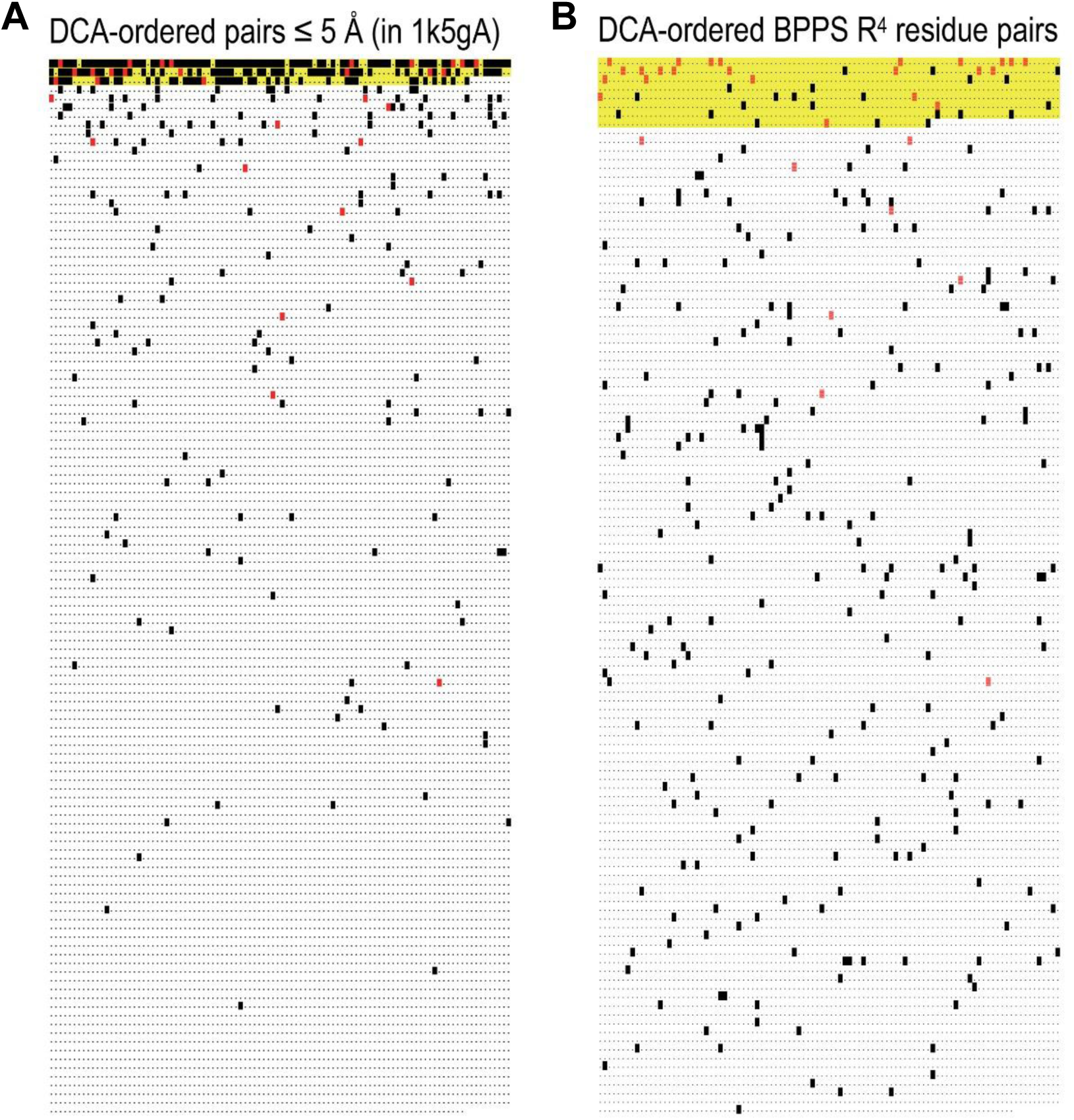
Distinguished residue pairs within an array of length *L* = 12,090, ordered by DCA scores for Ran GTPase (pdb: 1k5g), as computed by CCMpred using the R^4^ MSA (see Table 3). Distinguished pairs are represented by black and red blocks, the latter indicating pairs common to panels A and B; the remaining pairs are represented by dots. The region up to each cut point *X* is highlighted in yellow. **A**. Distinguished elements are those pairs separated by ≤ 5 Å in chain A of 1k5g. ICA results: *S* = 231; *D* = 354; *X*=291; *d* = 188; 53% of the distinguished pairs (*d*/*D*) occur in the initial 2.4% of the array (*X* /*L*). **B**. Distinguished elements are pairs of the 25 residues found by the BPPS program to be most distinctive of R^4^ GTPases. ICA results: *S* = 6.2; *D* = 281; *X* = 772; *d* = 46; 16% of the distinguished pairs occur in the initial 6.4% of the array. Note that because no ranking is available for the distinguished pairs in panel B we calculate *S* for both panels without the ball-in-urn component *P*_*b*_ and using only *P*_*a*_ [14].

**Figure S2.**
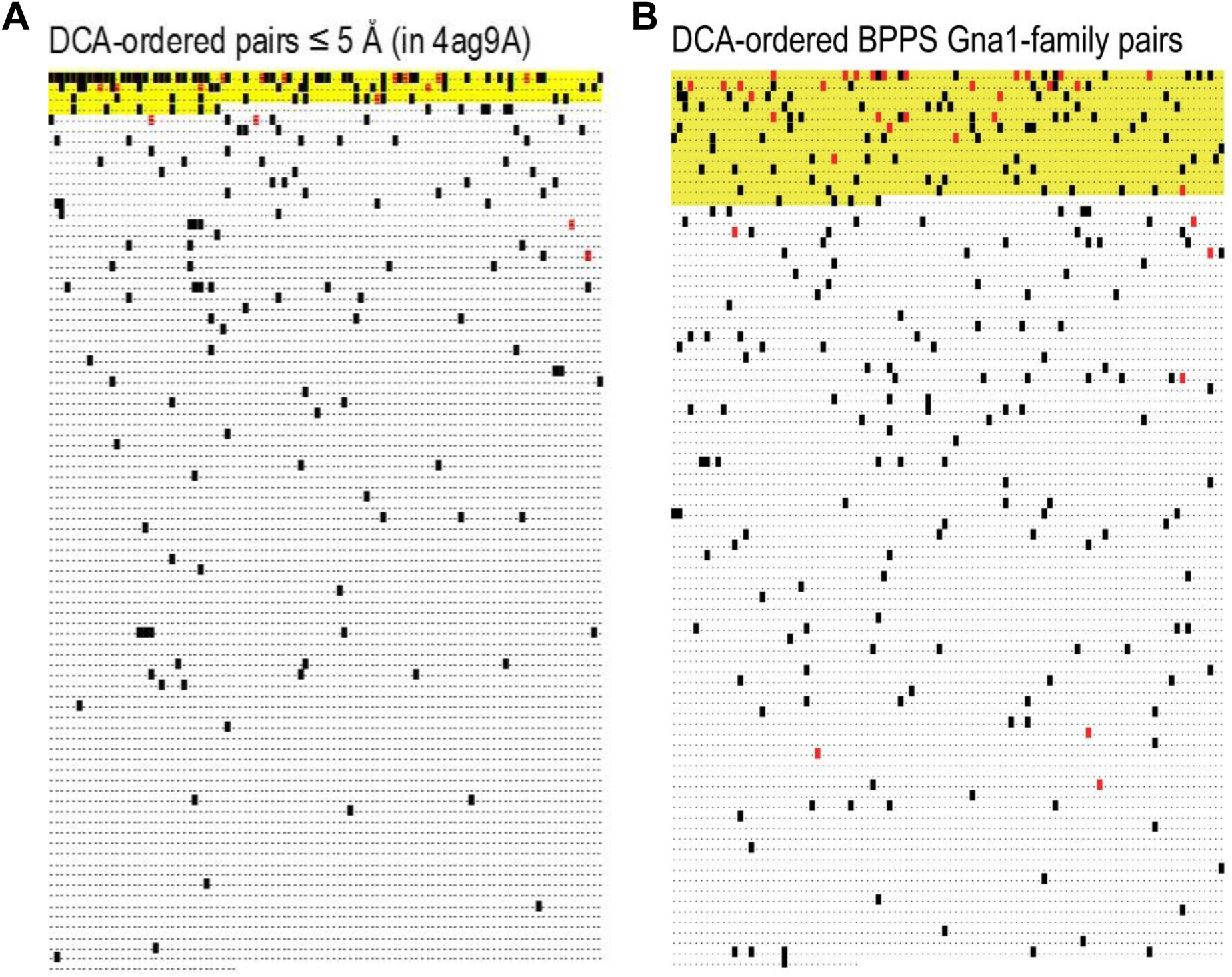
Distinguished residue pairs within an array of length *L* = 8,534, ordered by DCA scores for Gna1 (pdb: 4ag9), as computed by CCMpred using the MSA using for the analysis in Table 1. Distinguished pairs are represented by black and red blocks, the latter indicating pairs common to panels A and B; the remaining pairs are represented by dots. The region up to each cut point *X* is highlighted in yellow. **A**. Distinguished elements are those pairs in 4ag9 separated by ≤ 5 Å within chain A or between chains A and B (whichever is shorter). ICA results: *S* = 93; *D* = 271; *X*=331; *d* = 116; 43% of the distinguished pairs (*d*/*D*) occur in the initial 3.9% of the array (*X* /*L*). **B**. Distinguished elements are pairs of the 25 residues found by the BPPS program to be most distinctive of the Gna1 family. ICA results: *S* = 27.9; *D* = 263; *X* = 1238; *d* = 114; 43% of the distinguished pairs occur in the initial 14.5% of the array. Note that because no ranking is available for the distinguished pairs in panel B we calculate *S* for both panels without the ball-in-urn component *P*_*b*_ and using only *P*_*a*_ [14].

